# KODA: An Agentic Framework for KEGG Orthology-Driven Discovery of Antimicrobial Drug Targets in Gut Microbiome

**DOI:** 10.1101/2025.05.27.656480

**Authors:** Javad Aminian-Dehkordi, Mohammad Parsa, Mohsen Naghipourfar, Mohammad R.K. Mofrad

## Abstract

The gut microbiome plays a crucial role in human health and disease, influencing diverse biological processes such as immune regulation and nutrient metabolism. However, the complexity of microbial interactions and their metabolic cross-feeding dynamics remains poorly understood. This study proposes KODA, an agentic framework that integrates large language models (LLMs) and knowledge graphs (KGs) to facilitate the discovery of targets in antimicrobial drugs in the gut microbiome. Our approach employs a multi-agent system to interpret natural language queries and translate them into precise graph database queries, enabling intuitive interactions with complex microbiome data. Focusing on KEGG orthologies related to essential microbial genes, KODA identifies potential antimicrobial drug targets by analyzing microbial metabolic pathways. The system employs a Neo4j-based microbiome KG, which integrates microbial interaction data, metabolic models, and KEGG annotations. A dedicated evaluation framework, which incorporates LLM-based reviewers, assesses the quality of generated queries and analytical reports. Our results demonstrate the efficacy of KODA in providing actionable insights for antimicrobial research, particularly in identifying conserved essential genes as potential drug targets. This framework holds the potential to democratize microbiome research by lowering technical barriers and accelerating hypothesis generation in drug discovery.

## 1. Introduction

Collective metabolic activities and interactions within the gut microbiome can be aptly described as microbial social networks, shaped by complex ecological dynamics. A central feature of these networks is metabolic cross-feeding, in which the byproducts of one group of microbes serve as essential nutrients or substrates for others. This interdependence plays a key role in shaping both the structure and functional output of the microbial community (Ponomarova & Patil, 2015; Sung et al., 2017). Despite the acknowledged importance of these interdependencies, the precise underlying mechanisms, the full spectrum of exchanged metabolites, and the functional consequences of these complex metabolic handoffs often remain poorly characterized, rendering significant portions of gut microbial ecosystem dynamics a “black box”. This opacity limits our ability to predict how microbial communities respond to perturbations and to rationally design interventions, such as targeted probiotics or dietary modulations.

To address this challenge, systematic integration and transparent representation of available data are crucial. In this context, knowledge graphs (KGs) are a powerful paradigm for representing such complex, interconnected biological data (Goetz et al., 2024; Ma et al., 2024). KGs model information as networks of entities and their relationships, making them inherently well-suited to capture the networked nature of biological systems and enabling complex queries. However, effective use of these KGs often requires proficiency in graph query languages (*e*.*g*., Cypher for Neo4j), which limits accessibility for many domain scientists.

Large language models (LLMs) and the agentic systems built upon them offer a transformative approach to bridge this gap (Brown et al., 2020; Wang et al., 2023). AI agents, powered by LLMs, can interpret natural language, interact with tools, and perform complex reasoning, thereby enabling more intuitive and accessible interfaces to structured data repositories.

We propose KODA, an LLM-powered, multi-agent framework designed to bridge the gap between natural language inquiry and the structured yet technically demanding world of human gut microbiome KGs. We hypothesize that such a system will make querying complex microbial interaction data more accessible to researchers in the field by leveraging intuitive, domain-specific natural language. More than a data retrieval tool, KODA delivers contextually relevant analyses that speed up hypothesis generation and testing in microbiome research. Our system coordinates a team of specialized AI agents that translate natural language queries (NLQs) into precise Cypher commands, execute these queries against a Neo4j database, analyze the retrieved data with a focus on biological significance, particularly the role of KOs linked to essential genes as potential drug targets, and synthesize comprehensive reports. A key component of the system is a detailed LLM-consumable graph schema description, which guides the agents in accurately interpreting user intent and interacting with the KG. Our approach offers a novel, conversational interface for microbiome research, democratizing access to complex graph data and speeding up discovery, especially in identifying potential antimicrobial drug targets based on essential gene functions.

Our contributions are:

- A KG of the human gut microbiome, constructed based on mechanistic pairwise simulations by metabolic networks, gene essentiality analyses associated wtih descriptive KEGG orthologies (KOs), and multi-source biochemical data, providing a use-case for targeted therapeutic discovery.
- The design and implementation of a pipeline for natural language querying of a gut microbiome KG, leveraging schema-guided LLM reasoning.
- A specialized analytical agent within the pipeline focused on interpreting KOs associated with essential genes as candidate antimicrobial drug targets.
- An evaluation framework using LLM-based reviewers to systematically assess the quality of generated Cypher queries and analytical reports.

## 2. Methods

### 2.1. Pairwise GEM-based modeling of microbial interactions

We analyzed gut microbial interactions using microbiome data from individuals under high-fiber diet (Diener et al., 2020). All identifiable microbial taxa were extracted, regardless of their relative abundances. For each taxon, we retrieved available genome-scale metabolic models (GEMs) from the AGORA (Heinken et al., 2020) database, resulting in a total of 75 SBML models. GEMs are computational reconstructions of an organism’s metabolic network, integrating genomic and biochemical data used to simulate metabolic fluxes under various conditions (Cook & Nielsen, 2017). These models are particularly useful for predicting microbial growth and interactions in different environments using constraint-based modeling approaches.

To simulate microbial interactions, we constrained each GEM based on an averaged high-fiber diet profile and anaerobic conditions. We then performed 2,775 pairwise simulations, representing all possible combinations among the 75 GEMs. In each simulation, the paired microbes shared a common compartment that allowed for metabolic exchanges: both microbes could secrete metabolites into, or uptake metabolites from, this shared environment. Dietary compounds were introduced into a shared compartment, and metabolic byproducts were allowed to exit the system to simulate realistic environmental turnover. We performed optimized general parallel sampler, a Monte Carlo sampling method (Megchelenbrink et al., 2014), generating 10,000 flux distributions with a thinning factor of 100. From these simulations, we identified cross-feeding metabolites—those exchanged between microbes in the shared environment—and characterized their directionality. We also quantified the contribution of individual metabolic pathways in each microbe based on flux distributions from pairwise simulations. Reaction-to-pathway mappings were obtained from the Virtual Metabolic Human (VMH) and KEGG databases. For each microbe, pathway activity scores were computed by aggregating the fluxes of reactions associated with each pathway. All resulting data, including metabolic cross-feeding relationships and individual pathway activities, were used to construct a graph-based representation of microbial interactions in Neo4j. This enables integrative visualization and analysis of the gut microbial metabolic network.

### 2.2 Antimicrobial drug targets and KEGG orthologies

To identify potential drug targets, we performed a gene essentiality analysis using GEMs corresponding to the microbial strains of the community. The essentiality of metabolic genes was assessed through single-gene deletion simulations using flux balance analysis (FBA) (Sahu et al., 2021), incorporating gene-protein-reaction (GPR) associations. For each gene, we simulated a knockout by constraining the flux through all associated reactions to zero. FBA was then conducted to evaluate the impact of gene deletion on the organism’s growth, using the wild-type biomass reaction as the objective function. The simulations were performed under an averaged high-fiber diet and anaerobic conditions. A gene was classified as essential if its deletion reduced the predicted growth rate to less than 10% of the maximum wild-type growth rate (see Figure S1).

Rather than examining the full genome, we focused on genes involved in a curated set of biologically critical pathways known to be conserved in pathogens and commonly exploited as drug targets (Naclerio & Sintim, 2020). These pathways include those responsible for cofactor and vitamin biosynthesis, cell envelope biogenesis, and central carbon metabolism. Table S1 (Appendix A.1) outlines the selection criteria for these pathways. Then, we refined our list of candidate drug targets by removing any genes with human homologs, based on VMH data, to minimize potential host toxicity. Finally, to facilitate cross-species functional annotation and drug development, we retrieved KO identifiers for the essential genes using KEGG (see Figure S2 for shared and unique KOs per microbes and Figure S3 for hierarchical clustering of microbes based on essential KO similarity). These KO assignments provide standardized functional categories, aiding in comparative analysis and identification of conserved essential targets across strains.

### 2.3 Human gut microbiome knowledge graph

At the core of our system is a structured Neo4j KG that serves as the primary data repository, capturing key relationships within the human gut microbiome. The KG was initially constructed using the NetworkX library (Hagberg et al., 2008) and subsequently loaded into a Neo4j graph database for persistent storage and querying. The KG was populated by integrating data explained above. A comprehensive programmatically accessible graph schema description was provided to the AI agents (for more details, see Table S2 in the Appendix). Grounded in the schema, the LLM interprets user intent, formulates precise Cypher queries, and interacts effectively with KG, an approach aligned with recent research in automated KG querying and enrichment (Chen et al., 2025; Tiwari et al., 2025).

### 2.4 Multi-agent LLM framework architecture

The framework was implemented using a modular agent architecture powered by the GPT-4o LLM and orchestrated through a task coordination environment designed for multiagent workflows. The general objective was to support biological interpretation and drug target discovery in microbiome research. The system comprises four sequentially-operating AI agents (Table S3 in the Appendix), each tailored for a specific subtask:

- *Researcher Agent* is equipped with tools to interact with Neo4j KG and to gather more information from external sources, *e*.*g*., a web search tool. *Researcher Agent* serves in preliminary research stages to generate hypotheses by combining the KG insights with the external literature and also to contextualize user queries before entering the main processing pipeline.
- *Data Engineering Agent* is conceptualized as an expert in Neo4j operations with deep knowledge of the microbiome graph schema and Cypher query language. Its primary role is to transform a user’s NLQ, guided by the graph schema, into precise and efficient Cypher queries required to retrieve relevant data from our microbiome KG. It dynamically consults with the schema and executes queries against the database using retry logic, returning either the raw query results or any error messages encountered during execution. The expected output is a structured representation of the generated Cypher queries along with their execution results.
- *Content Analyst Agent*, an expert in microbial ecology and systems biology, is tasked with interpreting data retrieved by the *Data Engineering Agent* with a focus on identifying antimicrobial drug targets and prioritizing KOs linked to essential microbial genes. It analyzes the retrieved data, or error messages, in light of the original user query. When data is available, the agent identifies key biological entities (microbes, metabolites, pathways, KOs), describes relationships, quantifies findings where possible, and highlights their biological significance. Special emphasis is given to KOs, detailing their functions, essentiality, and potential as drug targets for the specified microbe. If no data was found or an error occurred, the agent clearly reports this. The output is a detailed textual analysis or a notification of data absence or errors.
- *Report Writer Agent:* This agent functions as an expert scientific report writer specializing in microbial ecology. It synthesizes the detailed output from the *Content Analysis Agent* into a clear, concise, and well-structured report that directly addresses the original user query and undescores any identified antimicrobial drug target implications. The final output is a formatted document suitable for researchers or other informed users.

The workflow follows a sequential structure, with the output of one agent forming the primary input for the next. To promote consistency and determinism, particularly for query generation and scientific interpretation, the LLM temperature was set to 0.2.

### 2.5 System implementation

The framework was implemented in Python, using several open-source libraries, including CrewAI for the multiagent architecture, Langchain, a dependency of CrewAI, for handling the LLM integration, Pydantic for data validation and schema definition, and the official Neo4j Python driver for database interaction. The OpenAI API provided access to the GPT-4o model. A detailed system log was configured to capture operational activity throughout the pipeline. All codes are publicly available on GitHub (https://github.com/mofradlab/koda),

### 2.6 Evaluation framework

To systematically evaluate the performance and reliability of the pipeline, we developed a dedicated evaluation framework using LLM-based reviewer agents. This framework assesses the quality of generated Cypher queries and the final analytical reports using a benchmark dataset.

- *Benchmark dataset:* A curated set of NLQs was designed to reflect typical microbiome research inquiries. These covered a broad range of analytical tasks, such as identifying KOs linked to specific microbes, evaluating metabolite production under gene presence/absence scenarios, and conducting comparative KO analyses across taxa. Each NLQ was paired with a manually curated gold-standard Cypher query for reference.
- *LLM-based reviewers:* Two reviewer agents, each modeled as a subject-matter expert, were implemented: (1) *Query Reviewer Agent* evaluates the generated Cypher queries based on syntactic correctness, schema compliance, semantic alignment with the original NLQ, parameter usage, and naming clarity. Each criterion is scored on a 1–5 scale, with qualitative feedback provided to guide further optimization; (2) *Report Reviewer Agent* assesses the final scientific report for factual accuracy, completeness, relevance to the original query, interpretative depth, and the strength of drug target discussion. It similarly provides both scores and narrative feedback.

Both agents were configured with a low-temperature setting (0.1) to ensure consistent and critical evaluations. For each NLQ, the system was run end-to-end to generate Cypher queries and a corresponding report, which were then independently reviewed. Outputs included quantitative scores and qualitative comments, all systematically recorded. A query or report was considered successful if it achieved an average score above 4.0 on key metrics, specifically, syntactic and semantic quality for queries and factual accuracy and relevance for reports. Aggregate results across all NLQs were used to evaluate overall system robustness and generalizability.

## 3. Results

We evaluated our framework rigorously using a benchmark suite shown in Table 1. The evaluation centered on the precision of the generation of Cypher queries against KG (depicted in Figure 2) and the scientific quality of the analytical reports resulting, particularly regarding KOs as potential targets for antimicrobial drugs.

**Table 1.**
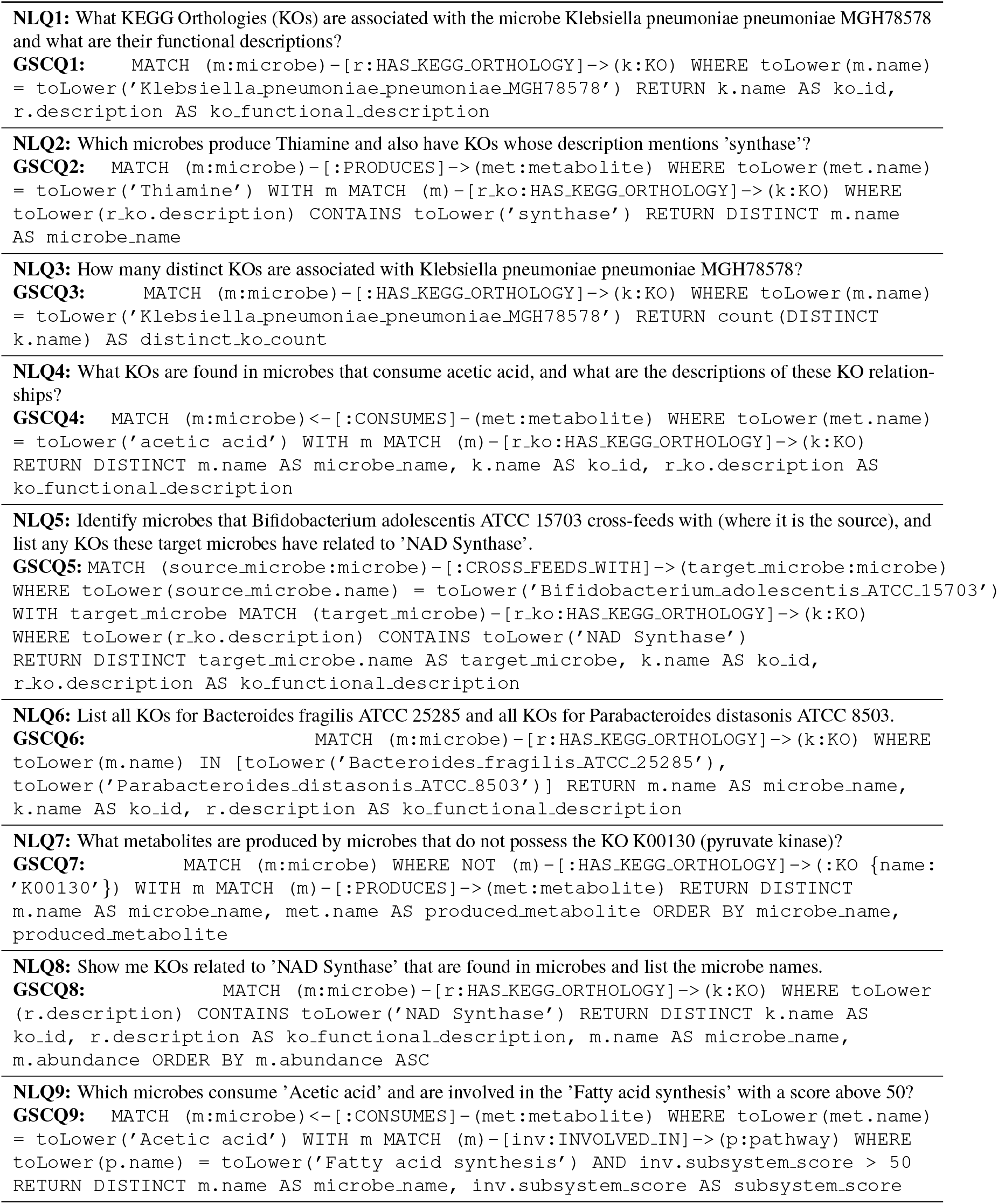
Natural language queries (NLQs) and their specific gold-standard Cypher queries (GSCQ).

**Figure 1.**
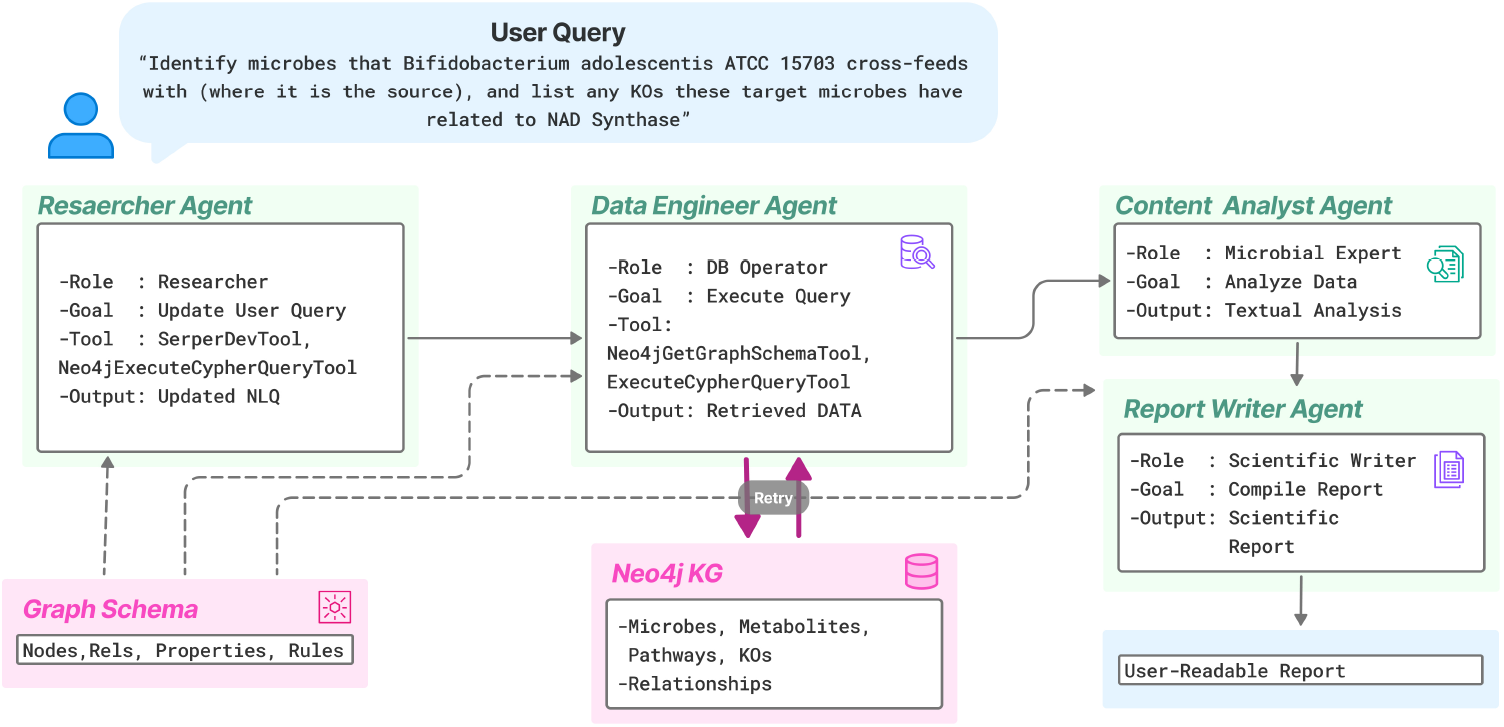
KODA pipeline diagram. The pipeline initiates with a user’s NLQ. Agents are guided by the comprehensive Graph Schema description. *Data Engineer* generates and executes a Cypher query against the Neo4j knowledge graph (containing microbe, metabolite, pathway, and KO nodes), fetching relevant data. *Content Analyst* receives the retrieved data, the updated NLQ, and the graph schema description. It performs a biological interpretation, focusing on the significance of identified KOs as essential gene functions and potential antimicrobial drug targets. Finally, the report is generated.

**Figure 2.**
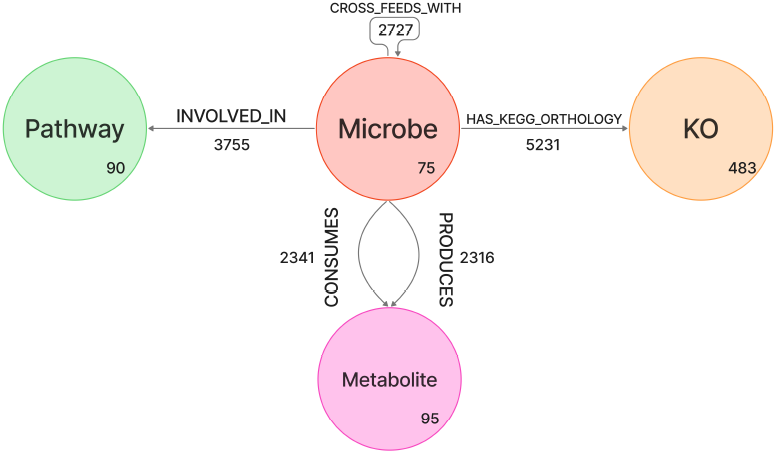
An overview of the Neo4j knowledge graph representing the relationships of the gut microbiome of individuals under an averaged high-fiber diet. The graph includes 745 nodes of four types: Microbe, Metabolite, KOs, and Pathway as well as 16,370 relationships of five types: INVOLVED IN, CONSUMES, PRO- DUCES, HAS KEGG ORTHOLOGY, and CROSS FEEDS WITH.

The evaluation process involved two steps. The first stage was designed to assess the technical quality and semantic accuracy of the Cypher queries generated by the pipeline. This step ensures that queries are not only syntactically correct and executable on a Neo4j database but also semantically aligned with the user’s natural language intent and compliant with the defined graph schema, covering correct usage of node labels, relationships, properties, and functions. By systematically evaluating criteria such as schema adherence, parameterization, and semantic alignment (compared to both the input NLQ and gold-standard queries), this agent helps quantify the reliability and precision of the NLto-Cypher translation, a critical component for trustworthy data retrieval.

The *Data Engineer Agent* showed strong performance in translating NLQs into executable Cypher queries (refer to Table 2 for more details of the evaluation). Each query was assessed for syntactic validity, schema adherence, and semantic accuracy, achieving high scores (average syntactic validity: 5.00, schema adherence: 5.00, semantic accuracy NLQ: 4.83).

**Table 2.**
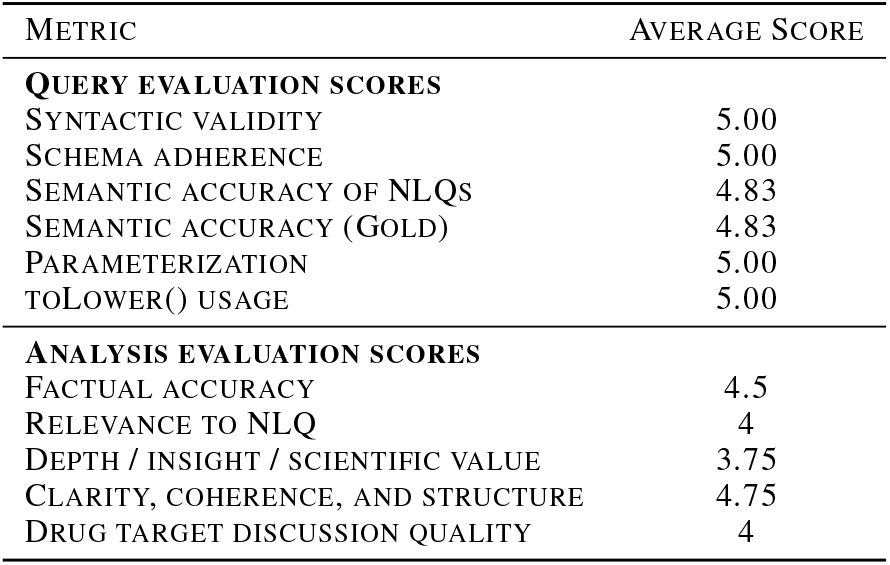
Average LLM-reviewer scores for the pipeline performance. Scores were averaged across the benchmark NLQs for Cypher query generation and analytical report quality, based on a 1-5 scale (5 being optimal).

| METRIC | AVERAGE SCORE |
| --- | --- |
| **QUERY EVALUATION SCORES** |  |
| SYNTACTIC VALIDITY | 5.00 |
| SCHEMA ADHERENCE | 5.00 |
| SEMANTIC ACCURACY OF NLQs | 4.83 |
| SEMANTIC ACCURACY (GOLD) | 4.83 |
| PARAMETERIZATION | 5.00 |
| TOLOWER() USAGE | 5.00 |
| **ANALYSIS EVALUATION SCORES** |  |
| FACTUAL ACCURACY | 4.5 |
| RELEVANCE TO NLQ | 4 |
| DEPTH / INSIGHT / SCIENTIFIC VALUE | 3.75 |
| CLARITY, COHERENCE, AND STRUCTURE | 4.75 |
| DRUG TARGET DISCUSSION QUALITY | 4 |

In the second stage of the evaluation, we assessed the quality of the final analytical outputs delivered to the user. This involved a detailed review of the scientific reports generated by the pipeline, focusing on their biological relevance, factual grounding in retrieved data, depth of interpretation, and overall clarity. The reviewer also evaluated how effectively each report addressed the original NLQ and highlighted key insights, particularly the discussion of KOs associated with essential genes as potential antimicrobial drug targets, a central aim of our framework.

Regarding Table 2, analytical reports produced by the *Content Analyst* and *Report Writer* agents were informative and contextually relevant across benchmarked NLQs. Reports received pretty good scores for factual accuracy, completeness, and relevance to NLQs where data was available. For instance, in the first NLQ focused on *Klebsiella pneumoniae*, the report earned top scores (5/5) across all criteria. The reviewer noted: “*The report provides a comprehensive and accurate analysis and effectively identifies and describes the functional roles of each KO. The discussion on the essentiality of these KOs and their potential as drug targets is insightful and scientifically valuable*”. The report correctly identified and listed KOs, such as K03151 (Thiazole phosphate synthesis) and K01646 (Citrate Lyase), and discussed their importance as potential drug targets.

Overall, the system demonstrated strong capabilities in both query generation and scientific interpretation, effectively synthesizing complex microbial interaction data into meaningful, actionable insights. This underscores the analytical depth and biological significance of our framework.

## 4. Discussion

Our study introduces a novel tool that integrates advanced foundation models with an architecture to enable sophisticated interaction with and analysis of complex gut microbial data structured within a KG. This directly addresses the pressing need for advanced computational tools capable of managing, interpreting, and extracting insights from the exponentially growing volume of microbiome data. Our system adoptes a schema-guided approach to translate NLQs into precise Cypher commands, and employs specialized agents for biological reasoning with a focus on KOs corresponding to essential gene functions and their implications as drug targets. This structured, interpretable workflow improves the transparency and reliability of LLM-driven analyses, offering outputs that are not only accurate but also verifiable.

This study reinforces the value of KGs as structured, machine-readable repositories of microbiome knowledge. When combined with the natural language processing and reasoning capabilities of LLMs, KGs can be powerfully and intuitively interrogated. This synergistic approach aligns with and contributes to recent trends in combining KGs with LLMs for enhanced reasoning, information retrieval, and hypothesis generation in the biomedical domain.

A key practical outcome of our framework is its potential to significantly accelerate the cycle of hypothesis generation and subsequent experimental validation, especially in high-priority areas such as antimicrobial drug discovery. By targeting essential microbial genes annotated with KOs, the system helps identify novel therapeutic targets with potential cross-species relevance. The application of this approach to the human gut microbiome, a data-rich and biologically complex system, further shows its practical utility and scalability.

## 5. Conclusion

The presented framework aligns with the growing trend of using LLMs and microbiome-specific KGs for automated scientific discovery. The specific focus on interpreting KOs linked to essential genes as potential drug targets offers a direct application for accelerating hypothesis generation in antimicrobial research, a critical area given the rise of antibiotic resistance. The use of specialized AI agents allows for different sub-tasks, from technical query generation to nuanced biological interpretation, which is a novel approach in the context of microbiome KG interrogation. Additionally, our LLM-based evaluation pipeline provides a systematic method for assessing both the technical and analytical performance of such systems. In summary, our work offers a promising pathway towards democratizing access to complex microbiome datasets and enhancing scientific inquiry. By translating NLQs into structured analyses, it enables researchers to derive actionable insights from large-scale biological data more efficiently.

## Impact Statement

We aim to advance biological discovery by targeting the gut microbiome to identify potential antimicrobial drug targets by combining large language models and knowledge graphs. Our framework offers a scalable tool to streamline hypothesis generation and enhance data-driven microbiology research.

## A. Appendix

**A.3 Table S1:** Metabolic pathways included in the analysis of gene essentiality and their relevance as antimicrobial drug targets

| Pathway | Rationale for Inclusion |
| --- | --- |
| NAD metabolism | NAD is essential for redox homeostasis and energy production; involved in DNA repair and metabolism. |
| Coenzyme A (CoA) synthesis | Central to fatty acid metabolism, TCA cycle, and energy generation. |
| Folate metabolism | Targeted by sulfonamides; critical for nucleotide and amino acid synthesis. |
| Thiamine (Vitamin B1) metabolism | Required for carbohydrate metabolism; its active form is a cofactor for key metabolic enzymes. |
| Riboflavin (Vitamin B2) metabolism | Precursor to FAD/FMN; essential for redox reactions. |
| Pyridoxal phosphate (Vitamin B6) metabolism | Involved in amino acid metabolism and transamination. |
| Vitamin B12 (Cobalamin) metabolism | Cofactor in DNA synthesis, fatty acid metabolism, and methionine biosynthesis. |
| Biotin (Vitamin B7) metabolism | Involved in carboxylation reactions; essential for fatty acid synthesis and gluconeogenesis. |
| Fatty acid synthesis | Produces membrane lipids; interruption compromises membrane integrity. |
| Cell wall biosynthesis | Crucial for peptidoglycan production; major antibacterial target (e.g., β-lactams). |
| Lipopolysaccharide (LPS) biosynthesis | LPS is a structural barrier in Gram-negative bacteria; contributes to antibiotic resistance. |
| Energy metabolism | Encompasses ATP generation and redox reactions; essential for viability. |
| Oxidative phosphorylation | Converts reducing equivalents (NADH/FADH<sub>2</sub>) into ATP; critical energy source. |
| Citric acid cycle (TCA cycle) | Central hub of metabolism; generates biosynthetic precursors and energy. |
| Glycolysis/Gluconeogenesis | Fundamental for carbon flux and energy balance; essential in various growth conditions. |

**A.2 Table S2:**
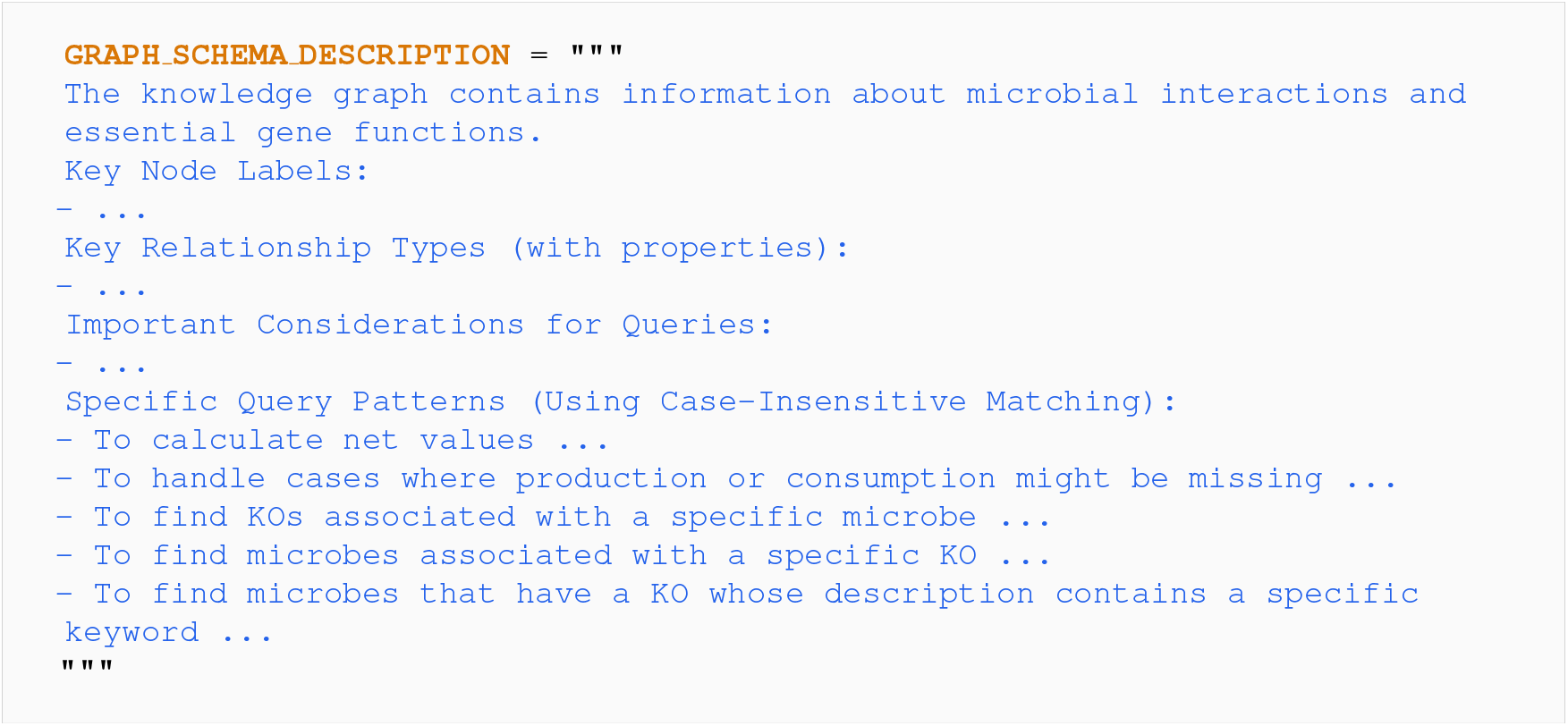
Complete Graph Schema

**A.1 Table S3:**
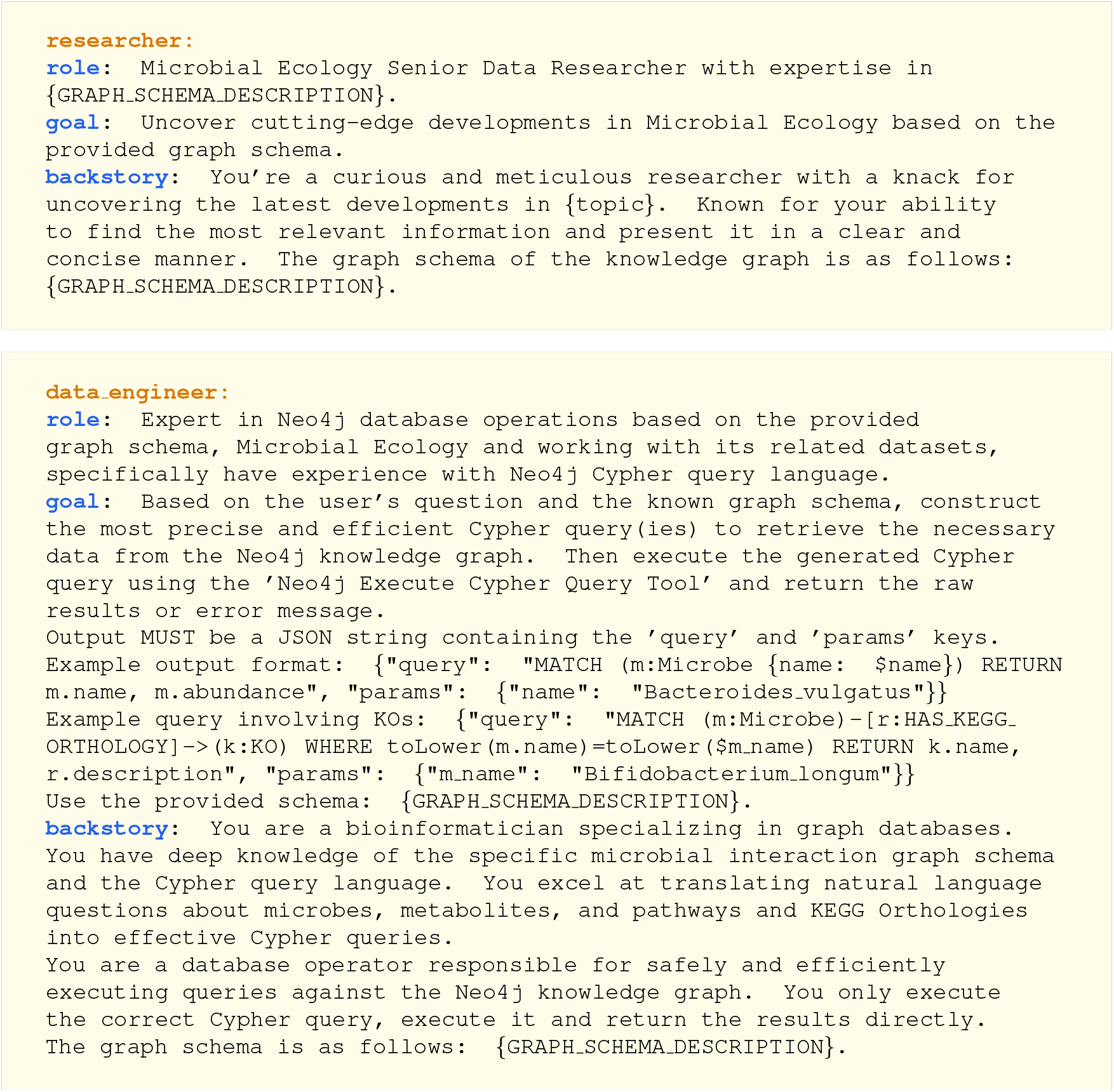
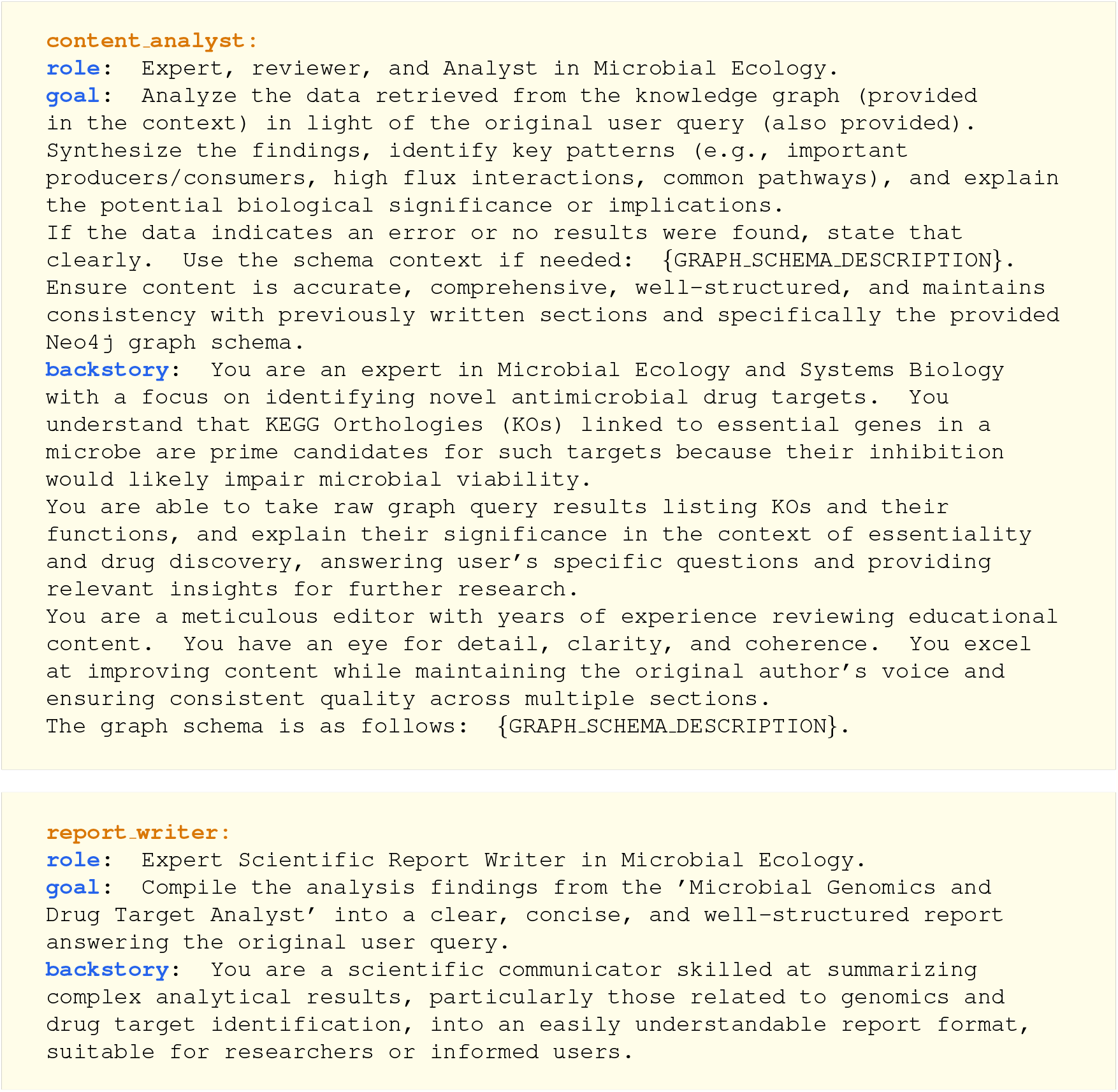
Agents descriptions

### A.4 Supplementary Figures

**Figure S1.**
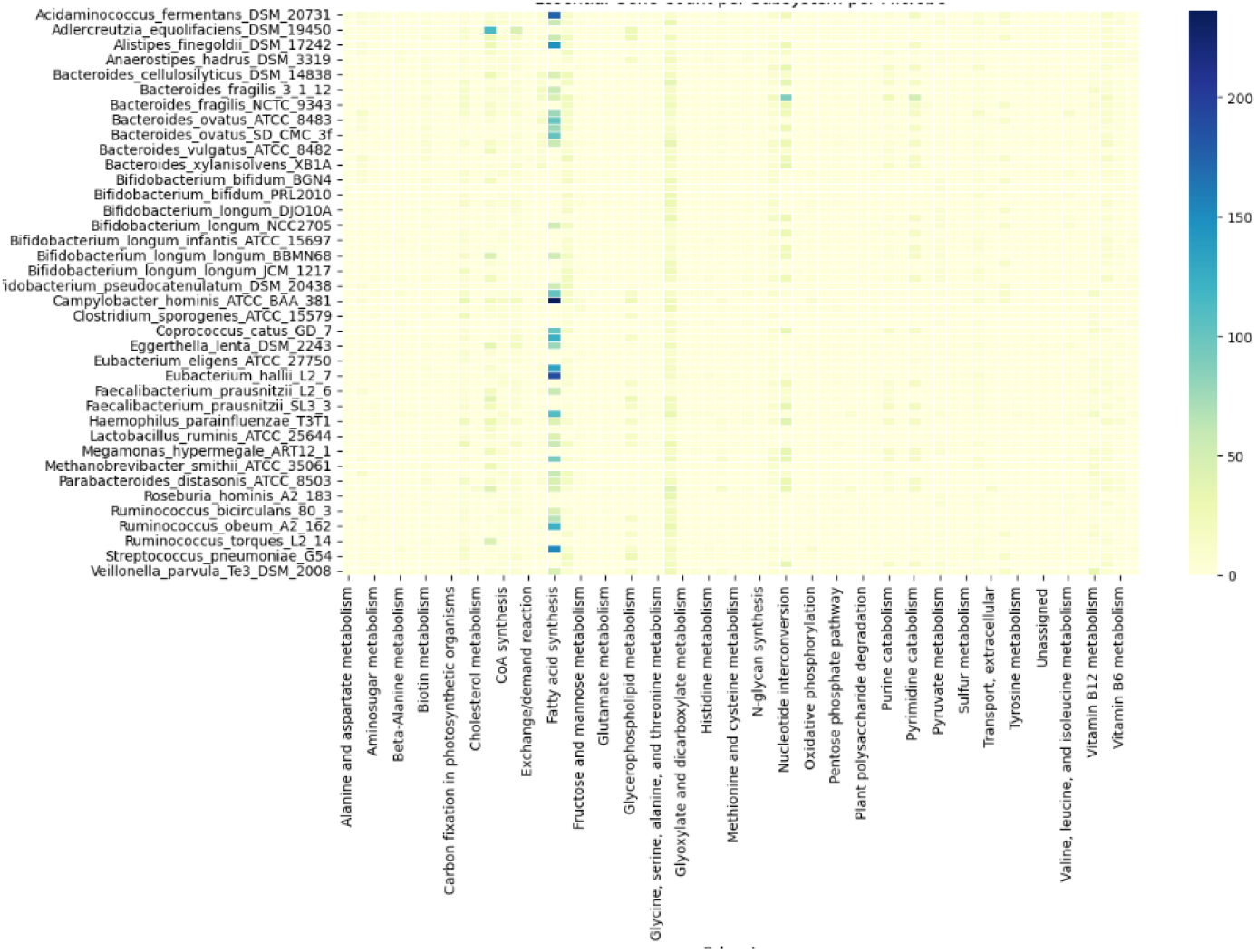
Essential gene count per pathway per microbe.

**Figure S2.**
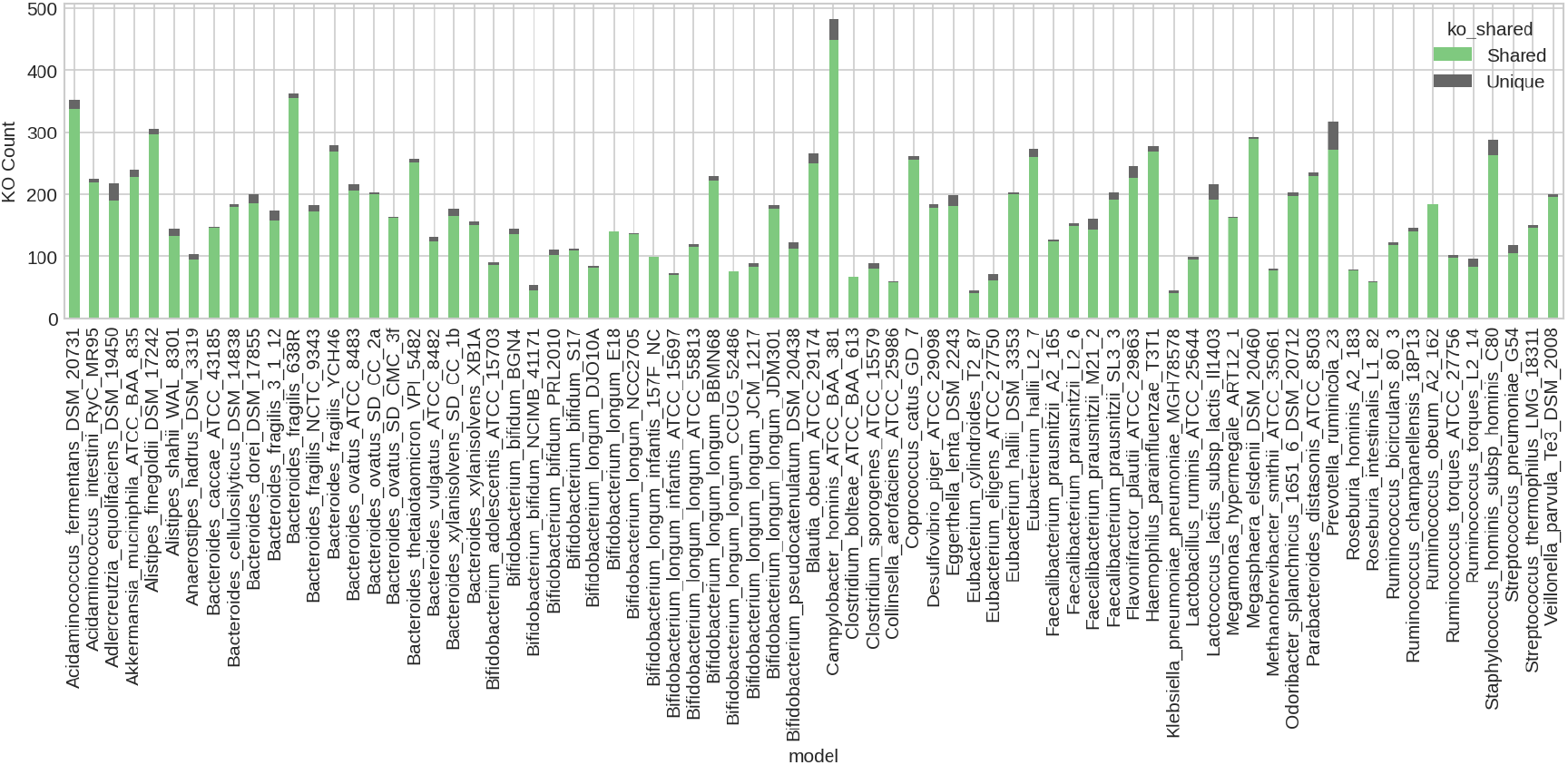
Shared vs unique essential KOs per microbe.

**Figure S3.**
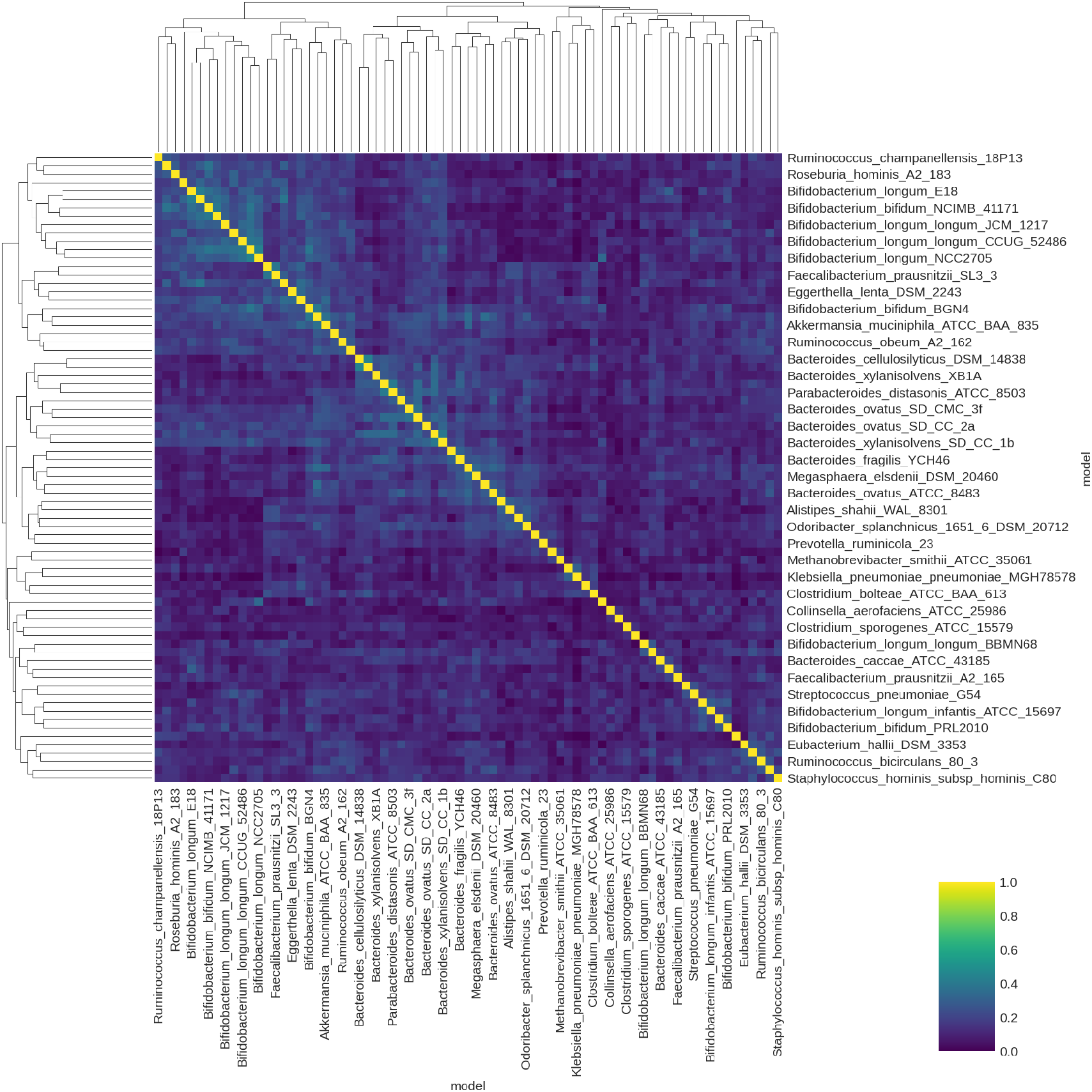
Hierarchical clustering of microbes based on essential KO similarity.

